# Contrasting patterns of genome-level diversity across distinct co-occurring bacterial populations

**DOI:** 10.1101/080168

**Authors:** Sarahi L Garcia, Sarah L R Stevens, Benjamin Crary, Manuel Martinez-Garcia, Ramunas Stepanauskas, Tanja Woyke, Susannah G Tringe, Siv G E Andersson, Stefan Bertilsson, Rex R. Malmstrom, Katherine D McMahon

## Abstract

To understand the forces driving differentiation and diversification in wild bacterial populations, we must be able to delineate and track ecologically relevant units through space and time. Mapping metagenomic sequences to reference genomes derived from the same environment can reveal genetic heterogeneity within populations, and in some cases, be used to identify boundaries between genetically similar, but ecologically distinct, populations. Here we examine population-level heterogeneity within abundant and ubiquitous freshwater bacterial groups such as the acI Actinobacteria and LD12 Alphaproteobacteria (the freshwater sister clade to the marine SAR11) using 33 single cell genomes and a 5-year metagenomic time series. The single cell genomes grouped into 15 monophyletic clusters (termed “tribes”) that share at least 97.9% 16S rRNA identity. Distinct populations were identified within most tribes based on the patterns of metagenomic read recruitments to single-cell genomes representing these tribes. Genetically distinct populations within tribes of the acI actinobacterial lineage living in the same lake had different seasonal abundance patterns, suggesting these populations were also ecologically distinct. In contrast, sympatric LD12 populations were less genetically differentiated. This suggests that within one lake, some freshwater lineages harbor genetically discrete (but still closely related) and ecologically distinct populations, while other lineages are composed of less differentiated populations with overlapping niches. Our results point at an interplay of evolutionary and ecological forces acting on these communities that can be observed in real time.

## Introduction

Bacteria represent a significant biomass component in almost all ecosystems and drive most biogeochemical cycles on Earth. Yet we know little about the population structure of bacteria in natural ecosystems and have yet to find and define the boundaries for ecological populations. Cohesive temporal dynamics and associations inferred from distribution patterns have been documented for many habitats and these observations are consistent with the notion of such populations as locally coexisting members of a species (Shapiro and Polz 2014). The most compelling cases are from collections of closely related isolates (Hanage et al 2005, Luo et al 2011, Shapiro and Polz 2014), but cultured species represent only a very small portion of the bacteria populating the Earth (Amann et al 1995, Hug et al 2016, Kaeberlein et al 2002), and thus we still know little about the most abundant lineages. Therefore it is critical to study microorganisms in their natural environments (Little et al 2008), in order to test if and how their population-level heterogeneity differs from the established models based on isolates. The advent of culture-independent approaches, such as single-cell genomics and metagenomics, provides an opportunity for gaining new insights about genome-level diversity at the population level for organisms that are currently difficult or impossible to culture.

The delineation of ecologically differentiated lineages within complex microbial communities remains controversial because direct evidence for such differentiation is usually sparse (Hunt et al 2008). Additionally, the appropriate level of phylogenetic resolution defining ecologically equivalent groups has not yet been established and likely varies across different groups (Fuhrman et al 2015). Past explorations for defining such groups have used genome-wide average nucleotide identity (gANI) across shared regions of isolate genome sequences (Konstantinidis and Tiedje 2005, Varghese et al 2015). These studies have found that gANI greater than 94-96% unites past classical species definitions and separates known sequenced strains into consistent and distinct groups. Genetically distinct populations have been identified in microbial communities using metagenomics by mapping reads against reference genomes and noting a coverage gap at 90-95% identity (Bendall et al 2016, Caro-Quintero and Konstantinidis 2012, Kashtan et al 2014, Konstantinidis and DeLong 2008, Oh et al 2011). Reads mapping with identities above the coverage discontinuity have been defined as originating from a ‘sequence discrete population’ (SDP) of genetically nearly identical cells that are distinct from other cells whose sequences map with identities below the coverage discontinuity (Bendall et al 2016). For the remainder of the manuscript, we will use the terms ‘population’ and ‘sequence-discrete population’ interchangeably.

We used a combination of time-series metagenomics and single cell genomics to define genetic diversification within ubiquitous and abundant freshwater lineages such as acI and tribe LD12. The term “tribe” was previously coined to delineate these groups using 16S rRNA gene sequences, where tribes are defined by monophyly and >97.9% within-clade 16S rRNA gene sequence identity (Newton et al 2007, Newton et al 2011). Freshwater microbial ecology researchers generally discuss and track these tribes as coherent units that are ecologically distinct from one another. A primary motivation for the present study was the challenge of moving beyond 16S rRNA sequence identity to delineate ecologically relevant taxonomic units given observed patterns of population-level heterogeneity, using shared genomic content. This study includes thirty-three Single Amplified Genomes (SAGs) representing fifteen phylogenetically coherent groups (i.e. freshwater “tribes”).

The SAGs in this study originated from four lakes geographically isolated from one another and represent a rich source of reference genomes that can be used to recruit metagenomic reads in order to study population-level heterogeneity and dynamics through time in naturally assembled communities. Two of the lineages featured in the present study are the abundant and ubiquitous freshwater Actinobacteria acI and Alphaproteobactera alfV containing the freshwater SAR11 sister-clade, tribe LD12. Members of these lineages are intriguing in their own right, as they represent groups of free-living ultramicrobacteria that dominate many freshwater ecosystems (Ghai et al 2014, Glöckner et al 2000, Heinrich et al 2013, Rösel et al 2012, Salcher et al 2010, Salcher et al 2011, Warnecke et al 2005, Zwart et al 2002). They differ markedly with respect to within-lineage diversity: LD12 is the sole tribe defined within the freshwater alfV lineage, while the acI lineage is comprised of 13 tribes (Newton et al 2011). The acI and alfV are not easy to cultivate in monocultures (Kang et al 2017) (though see (Henson et al unpublished data)) and share a large number of genomic and cellular traits. First, both lineages have genomes with GC content values lower than 40% and estimated sizes of about 1.5 Mb or less (Garcia et al 2013, Ghylin et al 2014, Kang et al 2017, Zaremba-Niedzwiedzka et al 2013). These genome characteristics are all the more striking since most cultivated species in the Alphaproteobacteria and Actinobacteria have GC-rich genomes up to 10 Mb in size. Second, both lineages have evolved by massive gene loss (Zaremba-Niedzwiedzka et al 2013). Third, the fraction of gained genes is only about 10% of the lost genes. Fourth, both groups of bacteria have small cell volumes (Heinrich et al 2013, Salcher et al 2011). However, acI and alfV seem to employ different substrate niche specialization. While acI is thought to primarily use polyamines, oligopeptides and carbohydrates, alfV specializes in carboxylic acids and lipids (Eiler et al 2016, Ghylin et al 2014, Salcher et al 2013).

By combining genome information from twenty-one previously published (Ghylin et al 2014, Zaremba-Niedzwiedzka et al 2013) and twelve new SAGs from different freshwater lineages and an extensive five-year time series of lake metagenomes (94 samples), we investigated the population-level heterogeneity of such ubiquitous freshwater bacteria for the first time. Our results confirm the existence of coherent sequence-discrete populations within these ubiquitous freshwater bacterial groups in natural communities and we could trace the abundance and gANI of these populations over monthly to seasonal time scales. Our work demonstrates the power of combining time-series metagenomics and single cell genomics for studying bacterial diversification and for describing ecologically meaningful population-level heterogeneity within communities inhabiting natural ecosystems.

## Results

### The SAG collection represents multiple clades within cosmopolitan freshwater lineages

We analyzed 33 SAGs from four different freshwater lakes. Twenty-one of these SAGs were previously analyzed for their genomic features and phylogenetic relationships (Eiler et al 2016, Garcia et al 2013, Ghylin et al 2014, Zaremba-Niedzwiedzka et al 2013). The 33 SAGs had total assembly sizes between 0.33 and 2.42 Mbp and were organized into 8 to 103 contigs with GC contents between 29.1% and 51.7% (Table 1). Estimated genome completeness, calculated using two different methods, ranged between 30% and 99%. Throughout the paper we will use mostly the shorter name version to facilitate reading, for example, M14 in place of AAA027-M14.

**Table 1.** Metadata for the 33 SAGs and genomes from (Kang et al 2017). The Genome OID is the object identifier for the genome record in the Joint Genome Institute’s IMG/MER Database. Estimated Genome Completeness was calculated using CheckM as described in the main text and (Parks et al 2015).

| Genome name | Genome OID in IMG/MER | Phylum/Class | Tribe | Lake | Collection date (M/D/Y) | Assembly size (Mb) | Est. Genome Comp. (checkm) | Number of contigs | GC content (%) | Citation |
| --- | --- | --- | --- | --- | --- | --- | --- | --- | --- | --- |
| AAA278-O22 | 2236661007 | Actinobacteria | acl-A1 | Damariscotta | 09/18/09 | 1.14 | 74.4 | 43 | 47.6 | Ghylin et al (2014) |
| AAA027-M14 | 2236661003 | Actinobacteria | acl-A1 | Mendota | 12/5/09 | 0.82 | 43.1 | 22 | 47.3 | Ghylin et al (2014) |
| IMCC25003 | 2602042019 | Actinobacteria | acl-A1 | Soyang | Jun-13 | 1.35 | NA | 1 | 49.1 | Kang et al (2017) |
| IMCC26103 | 2602042020 | Actinobacteria | acl-A4 | Soyang | Apr-14 | 1.46 | NA | 1 | 47.0 | Kang et al (2017) |
| AAA028-I14 | 2619618809 | Actinobacteria | acl-A6 | Mendota | 12/5/09 | 0.78 | 39.66 | 54 | 45.2 | This paper |
| AAA044-N04 | 2236661005 | Actinobacteria | acl-A7 | Damariscotta | 04/28/09 | 1.29 | 79.59 | 23 | 45.6 | Ghylin et al (2014) |
| AAA041-L13 | 2519899769 | Actinobacteria | acl-A7 | Damariscotta | 04/28/09 | 1.38 | 74.14 | 103 | 44.2 | This paper |
| AAA024-D14 | 2264265190 | Actinobacteria | acl-A7 | Sparkling | 05/28/09 | 0.78 | 48.4 | 82 | 45.4 | Ghylin et al (2014) |
| AAA023-J06 | 2236661001 | Actinobacteria | acl-A7 | Sparkling | 05/28/09 | 0.70 | 34.48 | 98 | 45.1 | Ghylin et al (2014) |
| IMCC19121 | 2602042021 | Actinobacteria | acl-A7 | Soyang | Oct-11 | 1.51 | NA | 1 | 45.5 | Kang et al (2017) |
| AB141-P03 | 2236876028 | Actinobacteria | acl-B1 | Stechlin | 05/25/10 | 0.66 | 45.98 | 66 | 40.8 | Ghylin et al (2014) |
| AAA278-I18 | 2236661006 | Actinobacteria | acl-B1 | Damariscotta | 09/18/09 | 0.94 | 63.73 | 54 | 41.4 | Ghylin et al (2014) |
| AAA028-A23 | 2236661004 | Actinobacteria | acl-B1 | Mendota | 12/5/09 | 0.83 | 57.56 | 64 | 41.5 | Ghylin et al (2014) |
| AAA027-L06 | 2505679121 | Actinobacteria | acl-B1 | Mendota | 12/5/09 | 1.16 | 76.59 | 75 | 41.7 | Garcia et al (2013) |
| AAA027-J17 | 2236661002 | Actinobacteria | acl-B1 | Mendota | 12/5/09 | 0.97 | 65.26 | 81 | 42.1 | Ghylin et al (2014) |
| AAA023-D18 | 2236661009 | Actinobacteria | acl-B1 | Sparkling | 05/28/09 | 0.75 | 44.22 | 67 | 39.6 | Ghylin et al (2014) |
| AAA044-D11 | 2619618811 | Actinobacteria | acl-B4 | Damariscotta | 04/28/09 | 1.15 | 66.18 | 30 | 44.2 | This paper |
| IMCC26077 | 2606217181 | Actinobacteria | acl-C1 | Soyang | Apr-14 | 1.55 | NA | 1 | 51.3 | Kang et al (2017) |
| AAA027-D23 | 2524023172 | Actinobacteria | acSTL-A1 | Mendota | 12/5/09 | 0.94 | 44.01 | 18 | 48.0 | This paper |
| AAA028-N15 | 2619618810 | Actinobacteria | acTH1-A1 | Mendota | 12/5/09 | 0.83 | 45.98 | 19 | 38.0 | This paper |
| AAA487-M09 | 2236347068 | Alphaproteobacteria | LD12 | Damariscotta | 09/18/09 | 0.63 | 53.15 | 97 | 29.1 | Zaremba-Niedzwiedzka et al (2013) |
| AAA280-P20 | 2236876029 | Alphaproteobacteria | LD12 | Damariscotta | 09/18/09 | 0.72 | 65.06 | 65 | 29.6 | Zaremba-Niedzwiedzka et al (2013) |
| AAA280-B11 | 2236876032 | Alphaproteobacteria | LD12 | Damariscotta | 09/18/09 | 0.67 | 51.24 | 47 | 29.8 | Zaremba-Niedzwiedzka et al (2013) |
| AAA028-D10 | 2236347069 | Alphaproteobacteria | LD12 | Mendota | 12/5/09 | 0.93 | 81.64 | 57 | 29.6 | Zaremba-Niedzwiedzka et al (2013) |
| AAA028-C07 | 2236661008 | Alphaproteobacteria | LD12 | Mendota | 12/5/09 | 0.85 | 74.12 | 32 | 29.5 | Zaremba-Niedzwiedzka et al (2013) |
| AAA027-L15 | 2236876031 | Alphaproteobacteria | LD12 | Mendota | 12/5/09 | 0.72 | 68.68 | 56 | 29.4 | Zaremba-Niedzwiedzka et al (2013) |
| AAA027-J10 | 2236876030 | Alphaproteobacteria | LD12 | Mendota | 12/5/09 | 0.79 | 69.56 | 82 | 29.8 | Zaremba-Niedzwiedzka et al (2013) |
| AAA027-C06 | 2264265094 | Alphaproteobacteria | LD12 | Mendota | 12/5/09 | 0.78 | 82.29 | 90 | 29.6 | Zaremba-Niedzwiedzka et al (2013) |
| AAA024-N17 | 2236876027 | Alphaproteobacteria | LD12 | Sparkling | 05/28/09 | 0.33 | 30.19 | 45 | 30.1 | Zaremba-Niedzwiedzka et al (2013) |
| AAA023-L09 | 2236661000 | Alphaproteobacteria | LD12 | Sparkling | 05/28/09 | 0.77 | 68.1 | 76 | 29.4 | Zaremba-Niedzwiedzka et al (2013) |
| AAA027-G08 | 2619618806 | Bacteroidetes | bacl-A1 | Mendota | 12/5/09 | 1.32 | 59.36 | 36 | 35.5 | This paper |
| AAA027-N21 | 2619618807 | Bacteroidetes | Flavo-A2 | Mendota | 12/5/09 | 2.21 | 92.44 | 36 | 33.1 | This paper |
| AAA027-K21 | 2619618803 | Betaproteobacteria | betIII-A1 | Mendota | 12/5/09 | 1.38 | 42.1 | 21 | 51.5 | This paper |
| AAA028-K02 | 2619618804 | Betaproteobacteria | LD28 | Mendota | 12/5/09 | 0.56 | 34.48 | 8 | 37.5 | This paper |
| AAA027-I06 | 2619618802 | Betaproteobacteria | Lhab-A1 | Mendota | 12/5/09 | 1.52 | 39.38 | 79 | 50.9 | This paper |
| AAA027-C02 | 2619618801 | Betaproteobacteria | PnecC | Mendota | 12/5/09 | 1.27 | 61.93 | 49 | 43.7 | This paper |
| AAA027-I19 | 2619618805 | Verrucomicrobia | Opiputaceae | Mendota | 12/5/09 | 2.42 | 54.58 | 63 | 51.7 | This paper |

NA = Not applicable

The 33 SAGs in the study represent fifteen different previously defined freshwater “tribes” (that are each monophyletic and defined by >97.9% within-clade 16S rRNA gene sequence identity, measured across the nearly full-length 16S rRNA gene) (Newton et al 2007, Newton et al 2011). Ten tribes are represented by only one SAG each, while four tribes (LD12, acI-A1, acI-A7 and acI-B1) have more than one SAG representative in our dataset. In addition to their classification based on 16S rRNA genes, the nine SAGs that were the only representatives of their lineage were classified using protein coding marker genes and PhyloSift (Darling et al 2014) (**Table S1**). To illustrate phylogenetic and taxonomic placement of the LD12 and acI SAGs, we used the PhyloPhlAn pipeline (Segata et al 2013) to generate a multi-gene tree (Figure 1A and 1B). The tree topology was consistent with previous phylogentic reconstructions for LD12 (Zaremba-Niedzwiedzka et al 2013) and acI (Ghylin et al 2014, Newton et al 2007). The tree supported the 16S rRNA gene-based tribe designations but did not reveal a clear biogeographic pattern, in agreement with previous analyses, i.e. members of the same tribes were found in different lakes (Zaremba-Niedzwiedzka et al 2013). However, our SAG collection was not designed to explore biogeography and much deeper sampling of each population would be needed to address this question rigorously.

**Figure 1.**
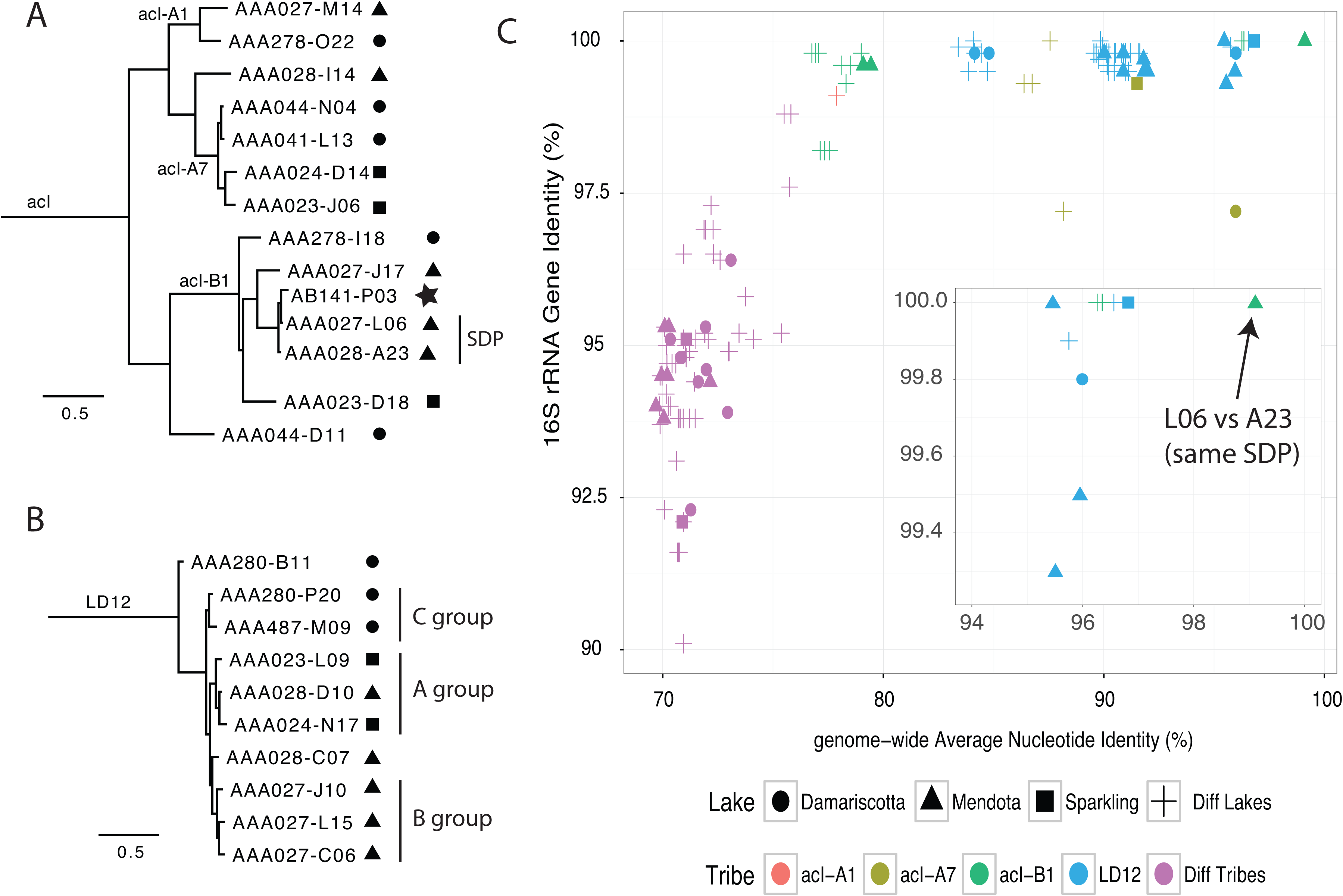
A. Phylogenetic tree of acI SAGs based on conserved single copy genes selected by PhyloPhlAn. Amino acid sequences from 400 genes were aligned. The tree topology is consistent with 16S rRNA gene-based phylogenies (Ghylin et al 2014). SAGs L06 and A23 are part of the same sequence discrete population (SDP) as defined in the text and further based on data shown in Figure 2. B. Phylogenetic tree of LD12 SAGs based on conserved single copy genes selected by PhyloPhlAn, representing 400 genes. The tree topology was consistent with prior work that provided evidence for finer-scale groups within the LD12 tribe (Zaremba-Niedzwiedzka et al 2013). C. Genome-wide nucleotide identity (gANI) versus 16S rRNA gene identity for pairs of SAGs. Alignment fractions for homologous genomic regions and 16S rRNA genes are given in **Table S2**. Shapes indicate the lake the tribe is from, if same, otherwise different lake is indicated. Colors indicate the tribe a pair is from, if same, otherwise different tribe is indicated. The arrow denotes the L06-A23 pair.

### Genome-wide nucleotide identity is consistent with phylogeny

Although multi-locus phylogenies supported the 16S rRNA gene based phylogeny, we wondered whether gANI could similarly be used to demarcate one tribe from another. To this end, we determined the pairwise gANI for genomes in the set of four tribes that each contained more than one SAG representative. This general approach has been proposed as a way to compare genome pairs using a single metric that robustly reflects phylogenetic and taxonomic groupings obtained using other polyphasic methods (Konstantinidis and Tiedje 2005, Varghese et al 2015). We asked whether all genome pairs from the same tribe shared a consistent minimum gANI. Most SAGs shared gANI of at least 78% and alignment fractions greater than 40% with other members of the same tribe (Figure 1C and **Table S2**). Most pairs from the same tribe that were also recovered from the same lake shared at least 84% gANI, but some pairs were much more similar (gANI above 95%). gANIs between pairs belonging to different tribes but still within the same lineage were markedly lower and typically below 74% (e.g. acI-A1 vs acI-B1) (Figure 1C and **Table S2**).

Although gANI is a useful univariate metric for comparing genome pairs, it masks the differences in sequence similarity of individual genes or genome regions that arise due to varying rates of divergence across loci. This variation can be visualized by plotting the frequency distribution of nucleotide identities calculated using a sliding window across the genome (Konstantinidis and Tiedje 2005). We asked whether different homologous genomic regions from two SAGs would have markedly different nucleotide identities even if they were from the same tribe. We used the most complete SAGs from the acI-B1 and LD12 tribes as reference genomes and calculated nucleotide identity using a sliding window with other SAGs from the same respective tribe and visualized the results as a frequency distribution (Figure 2 and **Figure S1**). The acI-B1 SAGs featuring the highest gANI (L06 and A23) were both from Lake Mendota and shared nucleotide identity consistently greater than 95% with a peak at 99-100%, suggesting they belong to the same SDP. The acI-B1 SAG P03 recovered from a lake in Germany had a frequency distribution with a peak more near 97% and a distinctly different shape. Other acI-B1 SAGs shared genomic regions with primarily 80-85% nucleotide identity. This was even true for J17, which was also collected from Lake Mendota and shared an average gANI of 79% with L06/A23 (**Table S2**), suggesting that cells belonging to the same tribe (acI-B1) and living in the same environment can have substantial genetic differences. The LD12 SAGs, which all belonged to the same tribe, also displayed three distinct patterns, with one peak near 85%, several near 91%, and two near 97%. Lake origin did not appear to explain these differences. That is, some LD12 cells from Lake Mendota were more similar to LD12 cells from Sparkling Lake than to other LD12 cells from Lake Mendota.

**Figure 2.**
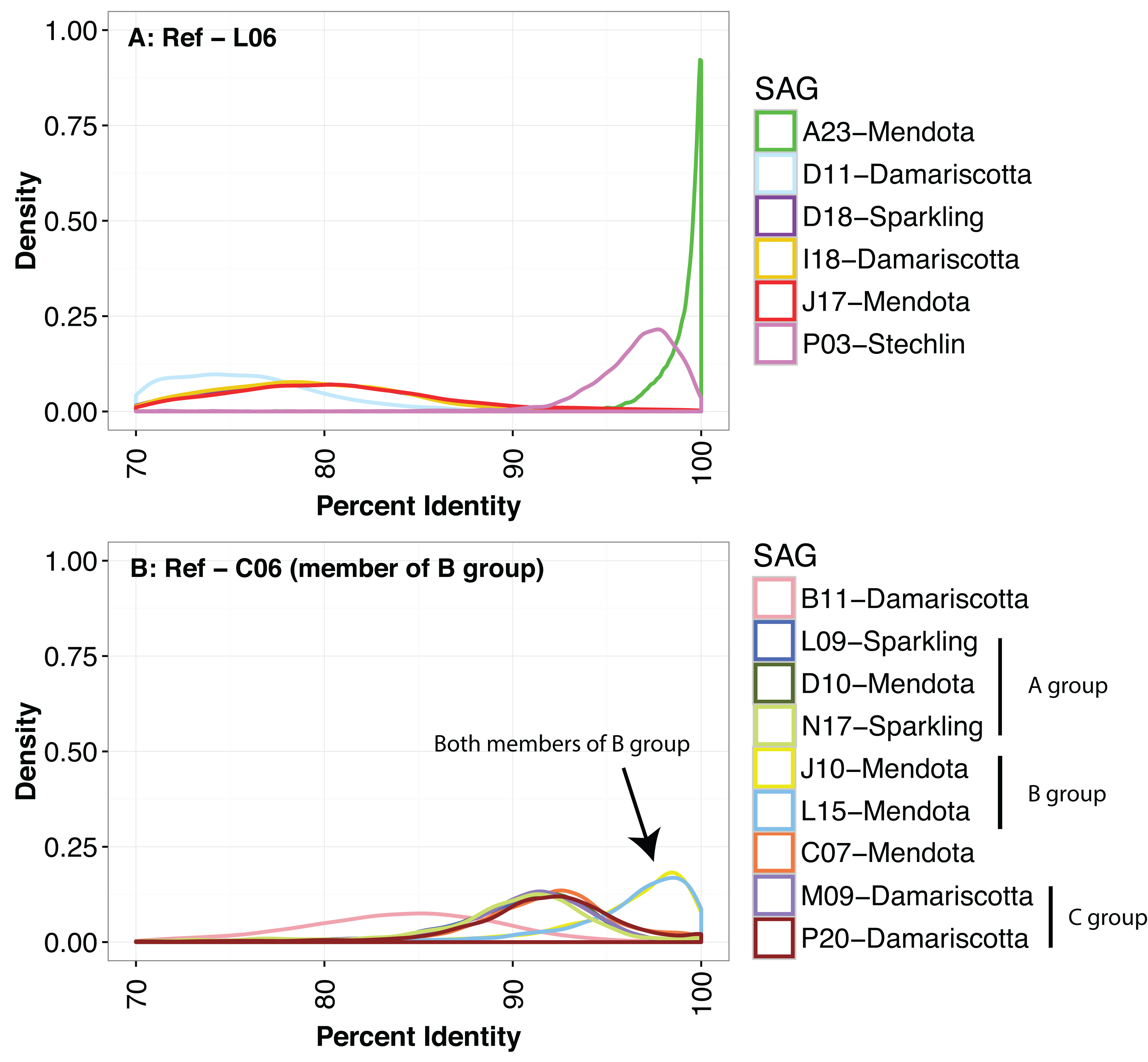
Nucleotide identity density plots for SAG versus SAG genome-wide comparison using a sliding window. Results are shown for two reference SAGs representing the most complete genomes from the most thoroughly sampled tribes. All SAG pairs were from the same tribe. Nucleotide identity was calculated with blastn using 301 bp fragments that overlapped by 150 bp. A. acI-B1 SAGs and other selected acI SAGs vs L06. Note that the purple line (D18) is hidden underneath the orange (I18) and red (J17) lines. B. selected LD12 SAGs vs C06. Note the dark blue line (L09) is hidden under the light green (N17) line. Group designations match those shown in Figure 1B, as proposed previously (Zaremba-Niedzwiedzka et al 2013). An expanded multi-panel version of the same data is shown in **Figure S1**, for clarity.

### Diversity of wild populations inferred using SAGs

The variety of patterns observed in Figure 2 indicated substantial within-tribe variability even among cells recovered from the same lake. This made us wonder if tribes were composed of genetically and ecologically distinct populations coexisting in the same environment. SAGs can serve as relevant reference points to study the diversity of abundant populations sampled using shotgun metagenomics by recruiting metagenomic reads and examining the extent of nucleotide identity for each aligned read (Stepanauskas 2012). The results can also be used to identify sequence-discrete populations whose boundaries are revealed by recruitment patterns and specifically the dramatic drop in coverage observed around 95% sequence identity (Bendall et al 2016, Caro-Quintero and Konstantinidis 2012, Konstantinidis and DeLong 2008). We asked whether such SDPs could be identified using metagenomic reads from Lake Mendota, WI, USA, by mapping them to the 33 SAGs, 19 of which were collected from this lake.

Each of the SAGs was first used to recruit reads from a single metagenomic dataset collected from Lake Mendota on 29 April 2009 (**Figure S2**). This time point was chosen because it was the sample collected closest to the date on which the single cells were collected (12 May 2009). Frequency distribution plots of the same data (Figure 3 and **Figure S3**) revealed patterns that were similar to those obtained with SAG pairs (Figure 2). The five acI-SAGs from Lake Mendota (J17, L06, A23, M14 and I14) recruited more reads than the acI-SAGs from other lakes, with many reads recruiting at nucleotide identity greater than 97.5% (Figure 3A). All of the acI-SAGs also recruited many reads at 60 – 90% identity (Figure 3A and D), creating the characteristic bimodal distribution observed in previous work (Caro-Quintero and Konstantinidis 2012). Based on these results, we hereafter consider reads sharing > 97.5% nucleotide identity as coming from the same, operationally defined *population* (i.e. SDP) as the reference SAG. Thus, the acI lineage in Lake Mendota on 29 April 2009 was composed of multiple SDPs. Interestingly, the acI-B1 tribe in Lake Mendota, a subset of the acI lineage, appeared to be composed of at least two coexisting and genetically distinct populations, one represented by SAG J17 and the other by SAGs A23 and L06, consistent with the pairwise gANI observed using only the SAGs (Figure 2).

**Figure 3.**
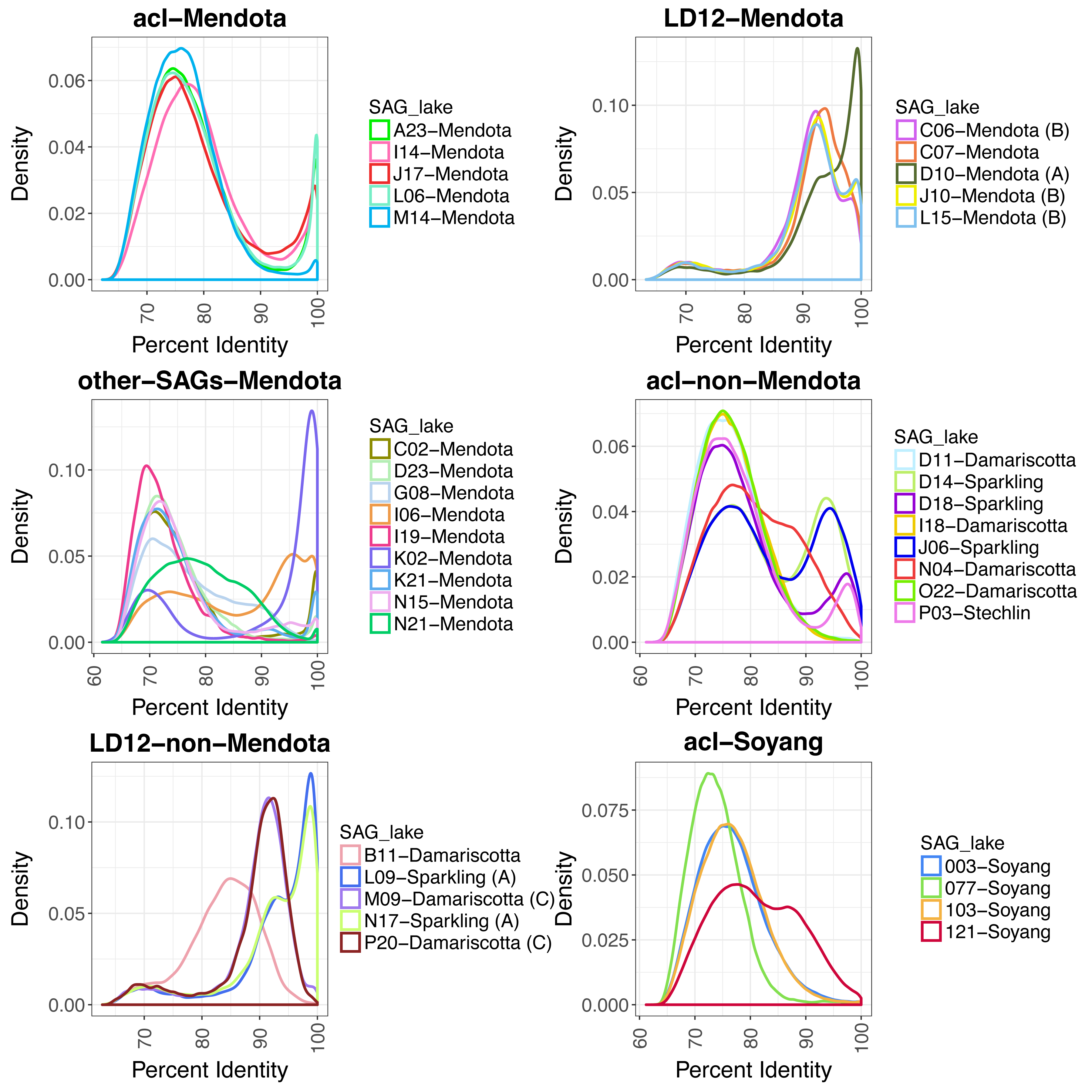
Mapping metagenomic reads from Lake Mendota to SAGs and four genomes from Lake Soyang (Kang et al 2017). The x-axis represents nucleotide identity of the recruited reads. The metagenome sample was collected from Lake Mendota on 29 April 2009. Reads were only counted if they aligned over a minimum of 200 bp. Recruitments were not competitive, meaning that each read could recruit to multiple SAGs. Analogous competitive recruitments that required each read to recruit to only one SAG are presented in **Figure S4**. The noncompetitive recruitment showed the close relationship of the LD12 populations that is not visible in the competitive recruitment. An expanded multi-panel version of the same data is presented in **Figure S3** for clarity. Each panel represents a different sub-set of the SAGs: A. acI from Mendota, B. acI not from Mendota, C. LD12 from Mendota (group members demarcated in legend), D. LD12 not from Mendota (group members demarcated in legend), E. other freshwater groups from Mendota, F. genomes from Lake Soyang, Korea. Regarding the other freshwater groups from Mendota, since each of these SAGs represent just one tribe, it is not appropriate to infer any general conclusions for these populations or tribes, but we present them here to show the intriguing diversity of recruitment patterns. We finally underscore the need to more deeply sample individual population members using SAGs, to better capture and describe the range of variation in population heterogeneity.

To determine if we recovered representative SAGs from all acI populations in Lake Mendota, we next performed recruitments competitively, allowing each read to only map to the SAG with the greatest percent identity (**Figure S4**). Since the patterns in Figure 3 were generated by non-competitive recruiting, some reads mapping with 100% identity to one SAG might for example also have mapped with 60-90% identity to SAGs from different SDPs. Under competitive recruiting conditions the resulting frequency distributions changed and the fraction of reads recruiting with 60-90% identity to each acI SAG dropped dramatically (**Figure S4**). However, a secondary peak around 80% identity still remained in most cases, and it is possible these reads originated from cells belonging to other acI populations lacking a representative SAG.

LD12 SAGs collected from Lake Mendota (C06, J10, L15, C07 and D10) also had a distinctive peak of recruited reads at >97.5% sequence identity (Figure 3B), although the overall shape of the recruitment patterns differed dramatically from those of the acI lineage. For example, LD12 SAGs had a secondary recruitment peak at ∼92% identity whereas the acI SAGs had secondary peaks at ∼75% with non-competitive recruiting (**Figure S4**). This suggests the SDPs within the LD12 tribe were more similar genetically than populations comprising the acI-B1 tribe. In fact, the populations were sufficiently similar that the hallmark coverage discontinuity below 97% similarity was not particularly pronounced (Figure 3B). Under competitive recruiting conditions, the LD12 recruitment distribution plots had remarkably different shapes (**Figure S4B and D**), as compared to the uncompetitive recruiting conditions (Figure 3B), and each SAG had only a single peak at >97.5% identity. This suggests the majority of LD12 cells in Lake Mendota belong to SDPs represented by the SAGs in our collection.

All but one (I06) of the other freshwater SAGs in this study that were collected from Lake Mendota generated the distinctive read recruitment frequency peak above 97.5% identity (Figure 3C) that was observed for acI (Figure 3A). A negligible number of reads recruited to the SAGs collected from other lakes under the competitive recruiting conditions (data not shown).

Four complete acI genomes recovered from Lake Soyang in Korea were recently published, and we included these in our recruitment analysis (Figure 3F). Three of the SAGs exhibited recruitment frequency distributions analogous to those obtained using acI SAGs from Sparkling Lake and Damariscotta Lake (Figure 3D), with very few reads mapping above 90% ANI. The distribution from one SAG (IMCC19121) was remarkably similar to that obtained from SAG N04, which was recovered from Damariscotta Lake in Maine. Both IMCC19121 and N04 are members of the acI-A7 tribe and share 89.8% ANI with each other.

### Are sequence-discrete populations within a tribe ecologically discrete too?

Results from a single metagenome sample suggested that individual tribes were composed of multiple genetically distinct populations that could be delineated and tracked using metagenomic read recruitment. Next we hypothesized that these populations might also be ecologically distinct and fill different realized niches. If so, we might expect these populations to display different temporal abundance patterns. We followed changes in population abundance through time by recruiting reads from a five-year metagenomic time-series applying a nucleotide identity cutoff of 97.5%, using only those SAGs derived from Lake Mendota. SAGs from the LD12 tribe recruited more reads than all of the acI SAGs summed together, on almost all sample dates (**Figure S5**).

Using the relative number of reads recruited as a proxy for abundance, we found the J17 population, which belonged to the acI-B1 tribe, to be the most abundant acI population in almost every sample (Figure 4A and 5A). The abundance of the J17 population was poorly correlated over time with the other acI-B1 population represented by L06 (maximum Spearman rank correlation = 0.256), indicating each population had a different temporal abundance pattern.

**Figure 4.**
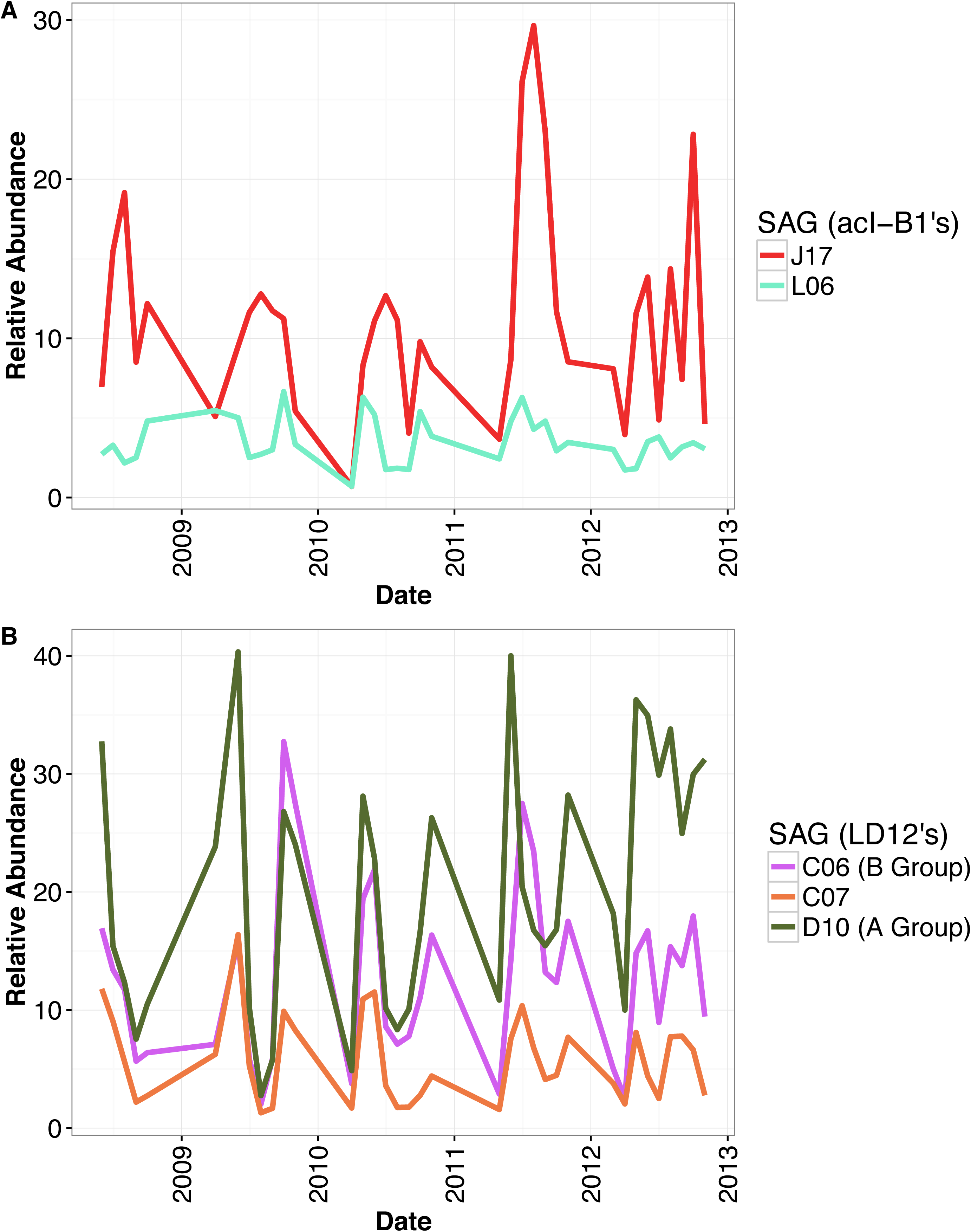
Sequence-discrete population abundance in Lake Mendota over time, as measured by the relative number of reads recruited to each SAG using blastn. All SAGs and samples are from Lake Mendota. Timepoints are pooled by month. Filtering criteria: ≥97.5% ANI and ≥200 bp alignment length. Recruitment was done using the most strict definition of competitive described in the methods, meaning any read that matched equally well to more than one SAG was not counted at all. Colors for each SAG are the same as in Figures 2 and 3. Relative abundance was calculated by normalizing the number of basepairs that recruited to each SAG by dividing by the genome size and the pooled metagenome size. The normalized number was then multiplied by the average pooled metagenome size. A. Relative abundance for each acI-B1 SAG. B. Relative abundance for each LD12 SAG. Membership in the groups defined in Figure 1B and by (Zaremba-Niedzwiedzka et al 2013) are denoted in the legend.

In contrast to the acI-B1 tribe, the populations comprising the LD12 tribe had highly similar abundance patterns. (Figure 4B and **S6**). The abundances of J10, L15, and C06 populations were strongly correlated (Spearman rank correlation = 0.996−0.999) (**Figure S8 and Table S5**) and tended to peak both in Spring and Fall (**Figure S6**). The D10 population was the most abundant in the dataset but its abundance was not as strongly correlated to the other LD12 populations (Spearman rank correlation = 0.705−0.725) (**Figure S8 and Table S5**). The C07 population was the least abundant but was also correlated to both the J10-L15-C06 populations and the D10 population (Spearman rank correlation = 0.850−0.873).

### Does the genetic diversity of populations change over time?

We also examined the extent to which within-population diversity varied through time by quantifying changes in population-wide ANI, i.e. the average identity of all reads mapping with at least 97.5% identity (Figure 5B). For this purpose, we only recruited reads to SAGs recovered from Lake Mendota. More abundant populations (such as LD12 and acI-B1 J17) generally had lower population-wide ANI variance through time compared to some less abundant populations (such as acSTL-A1-D23 and acI-A6-I14). For example, the SAG bacI-A1 G08 population had relatively high population-wide ANI in June 2009, around the time when the sample was collected for SAG library collection, but had markedly lower ANI on all other dates. One interesting exception to this observation was a significantly lower ANI for the relatively abundant acI-B1 L06-A23 population in 2012, as compared to 2007-2011 (Mann-Whitney U test p=1.4e-06).

**Figure 5.**
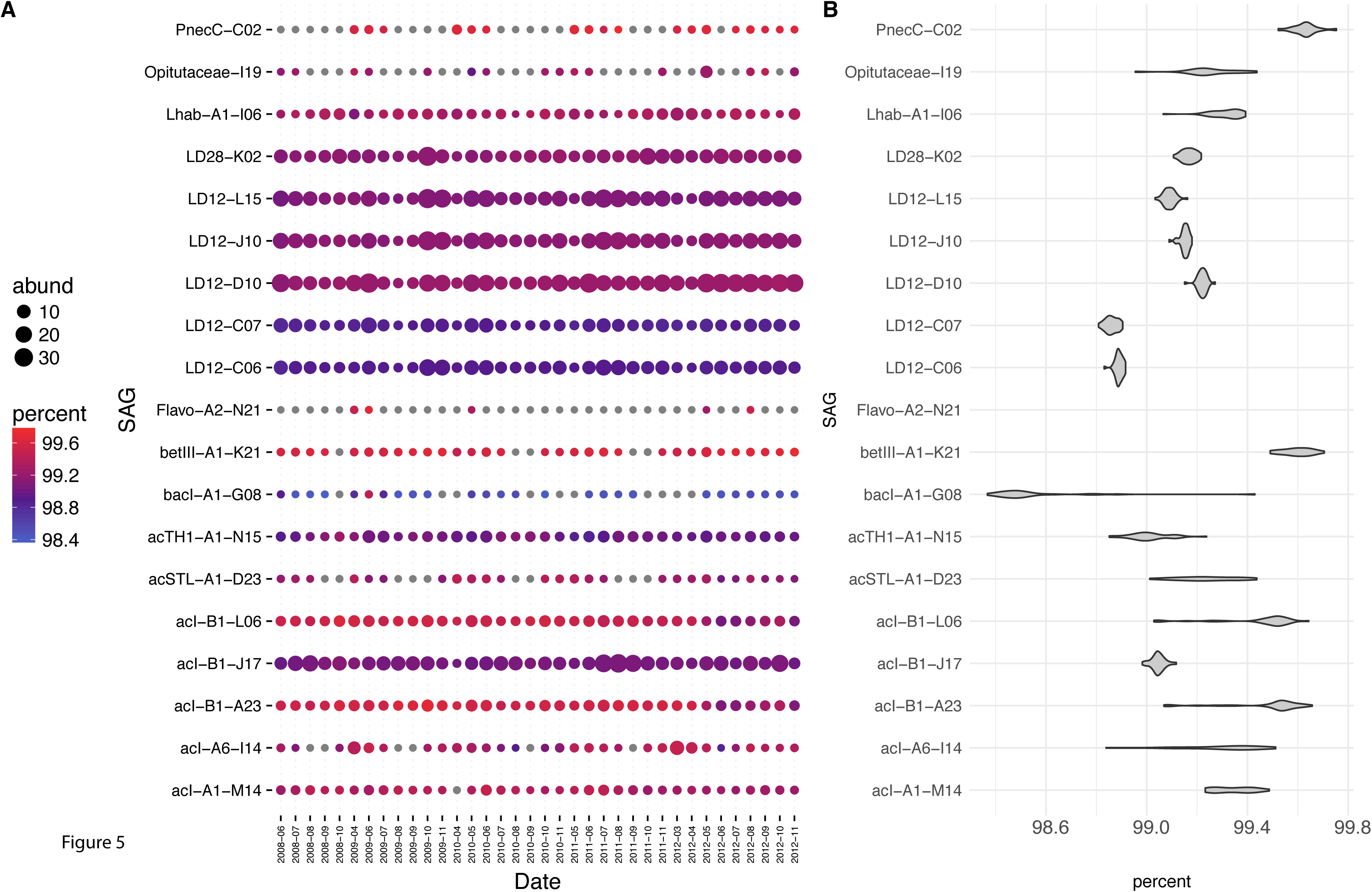
A. Metagenomic read recruitment using the SAGs from Lake Mendota. SAGs are in rows with bubbles representing all metagenomes from a particular month recruited against SAG. Filtering criteria: ≥97.5% ANI and ≥200 bp alignment length. Color scale indicates the ANI of the recruited metagenome reads. Bubble size represents the average coverage per base in the reference SAG divided by the size of the metagenome, multiplied by the average size of all metagenomes (1.34 Gigabases). Grey bubbles indicate that fewer than 200 reads recruited to the SAG in that month. Note that the resulting values do not represent a true measure of absolute abundance, but allow for quantitative comparison of month-to-month variation in population-level abundance. Recruitments were performed competitively, meaning that each read was counted for only one SAG, unless the read hit two SAGs equally well in which case it was counted for both SAGs. B. Variation in ANI for each SAG, across all 30 metagenomes from throughout the five years. Variation was not calculated for a SAG unless at least ten months recruited more than 200 hits each. The data underlying these plots can be found in Table S6.

## Discussion

Comparative genomics can reveal the diversity and structure of bacterial populations. This approach is particularly powerful when applied using single cells recovered from environmental samples (SAGs) and shotgun metagenomes from the same or similar ecosystems. Here we used a combination of 33 SAGs and 94 metagenomes collected over five years to ask the following questions: 1) How well do individual SAGs represent the population-level diversity found in natural communities? 2) Do common freshwater bacterial groups have similar patterns of population-level diversity? and 3) How stable is population abundance and diversity through time? We used the answers to these questions to gain insight into the population-level diversity and ecology of the cosmopolitan and abundant freshwater bacteria, alfV-LD12 (Alphaproteobacteria) and acI (Actinobacteria).

Sequence-discrete populations could be delineated in the Lake Mendota metagenome using our 33 SAGs as references, as has previously been demonstrated in other lakes using genomes assembled from metagenomes (Bendall et al 2016, Caro-Quintero and Konstantinidis 2012). We interpret the occurrence of these populations in the context of previously defined phylogenetically coherent and ostensibly ecologically distinct “tribes” composed of cells with >97.9% 16S rRNA identity (Newton et al 2011). We conclude that the freshwater tribes can contain multiple sequence-discrete populations. The converse is, of course, not true: sequence-discrete populations can never represent multiple tribes because tribes are by definition more distantly related to one another than genomes sharing a minimum of 97.5% gANI.

Pair-wise gANI analysis of SAGs and metagenomic read recruitment indicated that cells belonging to the same tribe but inhabiting different lakes were usually genetically distinct. For example, SAGs collected from other lakes generally recruited very few reads from Lake Mendota at ANI >97.5% while many recruited a substantial number of reads in the 89-92% range (Figure 3). However there were two prominent exceptions: LD12 N17 and L09, both of which are from Sparkling Lake. N17 and L09 share 97% gANI with Mendota SAG D10, which is substantially higher than the average (88%) and median (90%) within-tribe gANI (**Table S2**). These SAGs also recruited roughly the same number of reads with >97.5% identity as did the LD12 SAGs from Lake Mendota, though around 17% (L09) and 23% (N17) of the base pairs in the genomes did not recruit any reads. This implies that some gene content was present in the Sparkling Lake populations but missing in Lake Mendota. However, 10% of the base pairs in the D10 genome also did not recruit any reads, even though it was from Lake Mendota. We examined the phylogenetic distribution of low-coverage contigs and did not discern any evidence of contamination. This rare genome content could represent flexible or low frequency genes in the population, or contamination in the SAG preparation (Blainey 2013). However, it could also represent systematic coverage bias, a phenomenon that we are not able to rule out with the data at hand.

In Lake Mendota, acI cells are organized into genetically discrete populations, but the forces creating this organization remain unknown. The consistent lack of coverage around 90-97% identity in recruitment plots indicates Lake Mendota lacks acI genotypes sharing this degree of sequence similarity with our SAGs, or at least that these putative genotypes were consistently at much lower abundances than their close relatives over the five years surveyed. The P03 SAG from Stechlin Lake shares gANI of 96% with acI-B1 SAGs from Mendota, indicating that genotypes within this locally excluded sequence space do exist, at least as long as they are from different environments. We infer the persistence of the coverage discontinuity between populations to be less a factor of dispersal limitation and more likely the result of competitive exclusion and barriers to recombination within Mendota populations. Additional SAG and metagenomic studies are necessary to determine if similar coverage discontinuities are observed in other phylogenetic groups and in different environments. However, we do note that others have observed similar population-level diversity in other lakes (Bendall et al 2016, Caro-Quintero and Konstantinidis 2012) and marine ecosystems (Konstantinidis and DeLong 2008).

We know that both acI tribes and LD12 vary in abundance over seasonal and annual time-scales, based on previous work using 16S rRNA gene sequencing, quantitative PCR, and FISH (Allgaier and Grossart 2006, Eiler et al 2012, Heinrich et al 2013, Salcher et al 2011). Here we used our SAGs to track such populations at monthly intervals over five years (Figure 4 **and Figure S5**). The results confirmed prior work that showed acI tribes and LD12 are among the most abundant non-cyanobacterial groups in Lake Mendota (Newton et al 2011) but also revealed dynamics at unprecedentedly high phylogenetic resolution. Based on our extensive comparison of how SAGs recruited relative to one another, we are confident that our metagenomic recruitment filters allowed us to delineate discrete populations that would not be possible to resolve using more traditional and widely used methods (e.g. 16S rRNA gene sequencing or FISH). However, we do note that our acI SAG collection to date does not seem to fully capture the full diversity of acI populations in the lake, as evidenced by the residual peak of reads matching our SAGs at ∼70-80% ANI, even under competitive recruiting conditions. For example, we roughly estimate that our acI SAGs captured only 12% of the resident acI metagenome on 29 April 2009, as compared to 50% of the LD12 metagenome (**Table S3**). Thus, we cannot completely rule out the possibility that we missed strong correlations among other acI populations that we could not detect.

However, the most striking finding of our study was that metagenomic recruitments to LD12 SAGs yielded dramatically different patterns compared to the acI lineage. We discovered that LD12 populations were not as strongly genetically separated as acI populations; pair-wise gANIs between SAGs were higher and recruitment plots showed secondary peaks between 90-95% identity (Figure 3B), the same range where coverage of acI SAGs was at a minimum (Figure 3A). Under a competitive recruitment analysis, wherein each read is counted only once and attributed to the best match SAG, the secondary peaks disappear (**Figure S4**), indicating the LD12 SAGs represent highly similar, but still genetically discrete, populations. Temporal abundance patterns of these LD12 populations were strongly correlated over five years, whereas acI populations showed much lower correlation within tribes **(Figure S8**). This suggests that the acI-B1 populations are ecologically distinct (i.e. occupying temporally discrete niches) while LD12 populations are less differentiated with respect to niche dimensions, leading to co-occurrence and synchronization of temporal abundance patterns. LD12 is a particularly fascinating group because it is also a subclade of the broader SAR11 clade, with hypothesized ancient transition from marine to freshwater (Logares et al 2010) followed by specialization through gene flux and mutation, with comparatively low recombination rates (Zaremba-Niedzwiedzka et al 2013). Over time, low recombination rates and relatively low selection levels should lead to large genetic divergence among coexisting populations. Thus, we propose that LD12 populations are simply at earlier stages of differentiation as compared to acI populations, although we cannot exclude that something fundamental about their lifestyle is “holding” the populations together genetically and ecologically. This is particularly interesting in light of recent reports of unusually high recombination rates in LD12 (Zaremba-Niedzwiedzka et al 2013), pointing to the need to further investigate contrasting population structures and what these structures mean for the ecophysiology of the organisms. We do note that it is also possible that the highly correlated LD12 populations are each occupying unidentified distinct niches that are unrelated to the temporal correlation, allowing these slightly genetically differentiated populations to co-occur while being ecologically distinct. In any case, the lack of coherence among acI-B1 populations challenges our concept of tribes as ecologically coherent units and suggestions that freshwater microbial ecologists re-examine conventions for tracking these units through space and time. Taken together, these observations suggest fundamental differences in evolutionary history and/or lifestyles among these abundant and ubiquitous freshwater bacteria.

The metagenomic recruitments allowed us to also examine the extent to which diversity varied within and among populations as well as how diversity changed over time. We calculated the population-wide ANI for reads that recruited only above 97.5% and found the resulting value was remarkably stable through time for most of the abundant populations (Figure 5B). This was particularly true for the LD12 populations. However, one striking contrast was the acI-B1 population represented by L06/A23, which had consistent population-wide ANI of 99.3% during 2008-2011 but 99.0% during 2012 (Mann-Whitney U test p=1.4e-06). Similar shifts were observed previously in sequence-discrete populations inhabiting Trout Bog Lake, indicating this could be a common phenomenon among freshwater clades (Bendall et al 2016). Unlike the genome-wide selective sweep observed in one *Chlorobium* population from Trout Bog Lake, the distribution of single nucleotide polymorphisms within the L06/A23 population before and after 2012 exhibited no clear pattern of gene- or genome-wide sweep (data not shown). That is, it seems that the increase in population-wide gANI resulted in a change in the relative abundance of individual genotypes, rather than a single new genotype overtaking the population. It is difficult or impossible to separate genotypes within sequence-discrete populations using short-read shotgun sequencing, so further work using long-read technologies will be needed to link SNPs in populations to individual genomes. This kind of approach will likely be required to tease apart the paths leading to diversification within and among populations.

## Methods

### Single amplified genomes (SAGs)

Water samples (1-ml) were collected from the upper 0.5m to 1m of each of four lakes (Mendota, Sparkling, Damariscotta, Stechlin) and cryopreserved, as previously described (Garcia et al 2013, Martinez-Garcia et al 2011). These lakes were originally selected because they represent different freshwater trophic status (eutrophic, oligotrophic, mesoeutrophic, and oligotrophic, respectively) and geographic regions (Wisconsin and Maine, USA, and Germany). Bacterial single amplified genomes (SAGs) were generated by fluorescence-activated cell sorting (FACS) and multiple displacement amplification (MDA), and identified by PCR-sequencing of their 16S rRNA genes at the Bigelow Laboratory Single Cell Genomics Center (SCGC; http://scgc.bigelow.org). Thirty-two SAGs from lakes Mendota, Sparkling and Damariscotta were selected for sequencing based on the previously sequenced 16S rRNA gene as well as the kinetics of the MDA reactions (Martinez-Garcia et al 2011). The one SAG from Lake Stechlin was selected from a separate library because its 16S rRNA gene was 100% identical to an acI-B1 SAG previously analyzed (AAA027-L06) (Garcia et al 2013). In the present study we analyze 21 previously published and 12 new SAGs. All 33 SAGs were analyzed (Table 1) after genome sequencing, assembly, contamination removal and annotation as previously described (Ghylin et al 2014). Estimation of completeness was done using CheckM (Parks et al 2015) and the gene markers from a recent study examining a large collection of draft environmental genomes (Rinke et al 2013).

### Tree construction, Average Amino acid and Average Nucleotide Identity (AAI, ANI)

A phylogenomic analysis was conducted using PhyloPhlAn (Segata et al 2013). ANI was calculated by using the method described in (Konstantinidis and Tiedje 2005) with fragment size of 1000, minimum alignment length of 700 bp, percent identity of 70, and e-value of 0.001. AAI was calculated by averaging the identity of the reciprocal best hits from the BLASTP searches of the predicted proteins for each pair of genomes. 16S rRNA gene similarity for each pair was calculated using the overlapping region in an alignment created using a multiple alignment (default options) in Geneious Version R6 (Kearse et al 2012). Additional classifications were carried out using PhyloSift version 1.0.1, which examines 37 conserved single copy marker genes and places them into a phylogenetic reference tree (Darling et al 2014).

### SAG-to-SAG recruitments

SAG pairs from the same tribe were used to examine the frequency distribution of nucleotide identities across homologous regions of the two genomes. In order to create a sliding window for comparison, the contigs of all SAGs were shredded into 301bp fragments with 150 bp overlap. Two SAGs were selected as reference genomes: L06 as the most complete from the tribe acI-B1 and C06 as the most complete LD12. The contigs of each of the two selected SAGs were used as a reference for recruiting from the shredded SAGs using Blast 2.2.28 (Camacho et al 2009). Ribosomal RNAs were masked from the SAGs prior to performing blast.

### Five-year time series metagenome data: sampling, sequencing and recruitments

Samples were collected from Lake Mendota over the course of five years, as previously described (Kara et al 2013, Shade et al 2007). Lake Mendota, Madison, Wisconsin, (N 43°06, W 89°24) is one of the most well-studied lakes in the world, and is a Long Term Ecological Research site affiliated with the Center for Limnology at the University of Wisconsin Madison (Carpenter et al 2006). It is dimictic and eutrophic with an average depth of 12.8 m, maximum depth of 25.3 m, and total surface area of 3938 ha. Depth integrated water samples were collected from 0 to 12 m of the epilimnion (upper mixed layer) at 94 different time points during ice-free periods from summer 2008 to summer 2012, and filtered onto 0.2 μm pore-size polyethersulfone filters (Supor, Pall) prior to storage at −80°C. DNA was later purified from these filters using the FastDNA kit (MP Biomedicals). DNA sequencing was performed at the Department of Energy Joint Genome Institute using standard protocols (Walnut Creek, CA, USA). DNA from the 94 samples was used to generate libraries that were sequenced on the Illumina HiSeq 2000 platform. Paired-end sequences of 2 X 150bp were generated for all libraries. Adapter sequences, low quality reads (i.e. ≥80% of bases had quality scores <20), and reads dominated by short repeats of ≥3 bp were removed. The remaining high quality reads were merged with the Fast Length Adjustment of Short Reads (Magoc and Salzberg 2011) with a mismatch value of ≤0.25 and a minimum of ten overlapping bases from paired sequences, resulting in merged read lengths of 150 to 290 bp (**Table S4**). Metagenomes were pooled by month to reduce the time-series data to 30 observations and increase coverage. Original records can be found as a group in JGI’s Genome Portal: http://genome.jgi.doe.gov/Mendota_metaG.

All contigs from each of the 33 SAGs were used as a reference to recruit reads from the Mendota metagenomes using blastn. Metagenome reads that recruited to the SAGs were filtered and only alignments 200bp or longer were considered. An additional filter requiring an alignment percent identity of at least 97.5% was applied when analyzing the metagenome time series. Ribosomal RNAs were masked from the SAGs prior to performing the recruitments. Relative abundance was calculated by normalizing the number of basepairs that recruited to each SAG by the genome and pooled metagenome size and multiplying all by the average pooled metagenome size. When appropriate to the research question, recruitment was conducted “competitively”, meaning that if a read recruited to more than one SAG it was only counted for the best hit SAG. In this case, if a read recruited equally well to both SAGs, it was counted for both. In some cases we applied an even stricter definition of “competitive” and did not count any read that recruited equally well to more than one SAG. For Figure 3, recruitment was conducted “non-competitively”, meaning that reads could be counted for multiple SAGs as long as the hits met the filtering criteria. We note that this a commonly used approach developed by other researchers (Konstantinidis and Tiedje 2005, Konstantinidis and DeLong 2008). The figure and table legends contain the information necessary to discern which kind of recruitment criteria were applied for that specific analysis.

### Statistics, Visualization, Reproducible Methods

Datasets were analyzed and results were visualized using custom scripts written in R (R Core Team 2014) and python. Pipeline and scripts for analysis can be found at https://github.com/McMahonLab/blast2ani.

Supplementary information is available at the ISME Journal’s website.

## Acknowledgements

We thank Dr. Todd Miller and Sara Yeo for collecting the original water samples used to retrieve single cells from Lake Mendota and Sparkling Lake. We thank the Joint Genome Institute for supporting this work through the Community Science Program, performing the bioinformatics, and providing technical support. We thank Moritz Buck for informatics and statistical support. The work conducted by the U.S. Department of Energy Joint Genome Institute, a DOE Office of Science User Facility, is supported by the Office of Science of the U.S. Department of Energy under Contract No. DE-AC02-05CH11231. This research was performed using the compute resources and assistance of the UW-Madison Center for High Throughput Computing (CHTC) in the Department of Computer Sciences. The CHTC is supported by UW-Madison, the Advanced Computing Initiative, the Wisconsin Alumni Research Foundation, the Wisconsin Institutes for Discovery, and the National Science Foundation, and is an active member of the Open Science Grid, which is supported by the National Science Foundation and the U.S. Department of Energy’s Office of Science. KDM acknowledges funding from the United States National Science Foundation (NSF) Microbial Observatories program (MCB-0702395), the Long Term Ecological Research program (NTL-LTER DEB-0822700), an INSPIRE award (DEB-1344254), and the Swedish Wenner-Gren Foundation. RS acknowledges funding from NSF (DEB-0841933, EF-0633142 and OCE-821374). SB acknowledges funding from the Swedish Research Council. Sarahi Garcia thanks and acknowledges the JSMC for funding. MMG acknowledges funding from Ministry of Economy and Competitiveness (CGL2013-405064-R and SAF2013-49267-EXP)

### Author contribution

SLG, SLRS, RM, SB and KDM conceived the research. RM, MMG, TW and SGT conducted experiments and generated the data. SLG, SLRS and KDM analyzed the data. SLG, SLRS and BC prepared the figures. SLG, SLRS, RM, SB and KDM wrote the manuscript. All authors participated in revision of the manuscript.

### Additional Information

The raw shotgun metagenome reads and SAGs are publicly available in the JGI Genome Portal and via IMG/MER. The access number for each SAG and metagenome can be found in Table 1 and **Table S4**.

